# Constitutive 5-HT2C receptor knock-out facilitates fear extinction through altered activity of a dorsal raphe-bed nucleus of the stria terminalis pathway

**DOI:** 10.1101/2022.03.09.483588

**Authors:** Sandra T. Süß, Daniel Kremer, Linda M. Olbricht, Stefan Herlitze, Katharina Spoida

**Author notes:** Corresponding author: Dr. Katharina Spoida.

## Abstract

Serotonin 2C receptors (5-HT2CRs) are widely distributed throughout the brain and are strongly implicated in the pathophysiology of anxiety disorders such as post-traumatic stress disorder (PTSD). Although in recent years, a considerable amount of evidence supports 5-HT2CRs facilitating effect on anxiety behavior, the involvement in learned fear responses and fear extinction is rather unexplored. Here we used a 5-HT2CR knock-out mouse line (2CKO) to gain new insights into the involvement of 5-HT2CRs in the neuronal fear circuitry. Using a cued fear conditioning paradigm, our results revealed that global loss of 5-HT2CRs exclusively accelerates fear extinction, without affecting fear acquisition and fear expression. To investigate the neuronal substrates underlying the extinction enhancing effect, we mapped the immediate-early gene product cFos, a marker for neuronal activity, in the dorsal raphe nucleus (DRN), amygdala and bed nucleus of the stria terminalis (BNST). Surprisingly, besides extinction-associated changes, our results revealed alterations in neuronal activity even under basal home cage conditions in specific subregions of the DRN and the BNST in 2CKO mice. Neuronal activity in the dorsal BNST was shifted in an extinction-supporting direction due to 5-HT2CR knock-out. Finally, the assessment of DRN-BNST connectivity using antero- and retrograde tracing techniques uncovered a discrete serotonergic DRC-BNSTad pathway showing increased activity in 2CKO mice. Thus, our results provide new insights for the fear extinction network by revealing a specific serotonergic DRC-BNSTad pathway underlying a 5-HT2CR-sensitive mechanism with high significance in the treatment of PTSD.

## Introduction

Serotonin (5-hydroxytrytryptamine, 5-HT) plays a crucial role in the pathophysiology of diverse psychiatric disorders including schizophrenia, depression and anxiety disorders (1, 2). Depending on the receptor subtype involved, serotonin can induce both inhibitory and excitatory signals in neurons. The group of 5-HT receptors consists of seven families with at least 14 different receptor subtypes, making it challenging to define the explicit role of 5-HT in the respective disease (3). Especially the G-protein-coupled 5-HT2C receptor (5-HT2CR) has become a focus point of preclinical and clinical studies due to its close interaction with therapeutic drugs including serotonin reuptake inhibitors (SSRIs) (4, 5) Systemic administration of SSRIs is prevalently used to treat anxiety disorders and depression. However, initial drug intake often induces anxiogenic side effects associated with increased 5-HT2CR activation due to elevated 5-HT levels. In contrast, the beneficial long-term anxiolytic SSRI effect reflects desensitization of 5-HT2CRs (6). Anxiety disorders, in particular post-traumatic stress disorder (PTSD), are associated with exaggerated fear responses in combination with extinction deficits (7, 8). Under laboratory conditions, classical fear conditioning displays a common paradigm to study the neuronal correlates of learned fear and fear extinction in animal models (9, 10).

The amygdala is one of the key brain structures involved in the neuronal processing and storage of fear memories (9, 11–13). Particularly, a part of the extended amygdala, the bed nucleus of the stria terminalis (BNST), has gained attention in fear research over the past decade (14). Several lines of evidence have confirmed a BNST involvement in anxiety behavior. For instance, using optogenetic manipulation combined with behavioral tasks, Kim et al. (15) revealed that two subregions of the dorsal BNST exert opposite effects on the anxiety state. Activity of the oval nucleus (BNSTov) was associated with anxiogenic effects whereas activity in the anterodorsal subregion (BNSTad) was found to be anxiolytic. Additionally, two distinct studies using DREADD receptors (designer receptors exclusively activated by designer drugs) have confirmed the contribution of G-protein-coupled receptor (GPCR) signaling to the anxiety circuitry of the BNST (16, 17). Especially G_q_-pathway stimulation in GABAergic or 5-HT2CR-expressing neurons was reported to increase anxiety-like behavior. Similarly, activation of G_q_-coupled 5-HT2CRs in the BNST via direct infusion of the agonist meta-chlorophenylpiperazine (mCPP) produced an anxiety enhancing effect (16).

In contrast to the involvement of the BNST in anxiety behavior, its contribution to fear learning, especially to explicit cues and fear extinction is only poorly understood. Based on a model by Walker et al. (18), it has long been assumed that the BNST is involved primarily in fear learning to diffuse, unpredictable threats or contextual cues and the generation of sustained fear responses rather reflecting an anxiety-like state. In contrast, the generation of phasic fear in response to discrete threats or explicit cues was attributed to the central amygdala (CeA) (19–22). However, more recent work indicates the involvement of the BNST also in the processing of discrete, phasic cues (23–25).

Serotonergic fibers arising from the midbrain raphe nuclei, predominantly from the dorsal raphe nucleus (DRN), target the amygdala (26–28) as well as the BNST (29, 30). Aversive cues and stressors have been shown to increase 5-HT levels in the amygdala (31, 32). Similarly, aversive events as electrical foot shock presentation activate DRN neurons projecting to the BNST (17). Moreover, it has been described that anxiogenic drugs and negative stressors such as inescapable shocks activate 5-HT neurons in specific subregions of the DRN, including the dorsal subregion (DRD) and a caudal subregion (DRC) (33–35). However, the DRN represents a highly heterogeneous brain structure. Serotonergic neurons in the DRN are topographically organized, with anatomically and functionally distinct efferent and afferent projection neurons (36), but most studies investigating the DRN do not consider its high level of organization.

In the present study, we used an auditory fear conditioning paradigm in combination with mice constitutively lacking the 5-HT2CR to investigate the specific contribution of 5-HT2CRs to fear memory formation and fear extinction. In the first step, we used a fear conditioning and extinction protocol to verify behavioral differences between 5-HT2CR knock-out animals (2CKO) and WT littermates (WT). Our data revealed that exclusively fear extinction was facilitated in 2CKO mice. In a second step, we quantified the immediate-early gene product cFos, a commonly used marker for neuronal activity (37), after fear extinction in the DRN, amygdala and BNST. We found increased activity in the DRC subregion of the DRN in 2CKO animals even under basal home cage conditions. Significant alterations in cFos levels were also found in the BNST. In a last step, we used viral targeting of anterograde terminals and retrograde fluorogold tracing to uncover a discrete DRC-BNSTad projection, which may contribute to the extinction enhancing effect in 2CKO mice.

## Materials and Methods

### Subjects

Adult mice (9–16 weeks of age) were used for all experiments. Husbandry conditions included a 12 h light/dark cycle and constant room temperature. Food and water were freely available. All behavioral experiments include hemizygous 2CKO mice and WT littermates with a breeding background bearing C57BL6^htr2c+/htr2c-^ females (B6.129-Htr2c^tm1Jul^/J, stock no. 002627, Jackson Laboratory) and wildtype C57BL6 ^htr2c+/Y^ males (stock no. 000664, Jackson Laboratory). For anterograde labeling of neuronal terminals via double floxed tdTomato virus, Gad2-Ires-Cre mice (Gad2^tm2(cre)Zjh^/J, stock no. 010802, Jackson Laboratory) and ePet1-Cre mice (B6.Cg-Tg(Fev-cre)1Esd/J, stock no. 012712, Jackson Laboratory) were used. Mice were constantly group housed in cohorts of 3-5 individuals and isolated one week prior to behavioral experiments in a separate room next to the experimental room. All experiments were performed during the light phase. The experiments were approved by a local ethics committee (Bezirksamt Arnsberg) and the animal care committee of Nordrhein-Westfalen (LANUV; Landesamt für Umweltschutz, Naturschutz und Verbraucherschutz Nordrhein-Westfalen, Germany; AZ. 84-02.04.2016.A138). The studies were performed in accordance with the European Communities Council Directive of 2010 (2010/63/EU) for care of laboratory animals and supervised by the animal welfare commission of the Ruhr-University Bochum.

### Fear conditioning and extinction apparatus and procedure

Auditory fear conditioning and extinction were performed in a behavior cabinet (in-house production) consisting of a sound-attenuated chamber, provided with two speakers (Visaton FR 58, VISATON, Haan, Germany) for tone presentation, LED stripes (white and RGB) for context-dependent illumination and a camera (HD Pro C920, Logitech, Apples, Switzerland) for video recording (15 FPS). A fan enabled constant air circulation and background noise generation. The conditioning arena (25 cm x 25 cm x 37 cm) comprised exchangeable Perspex walls and several floors to generate different contexts. All behavior experiments include only male mice. The animals were habituated to handling for 4–5 days (5 min per animal each day) prior to the start of the behavioral experiments. Home cage control animals (HC) underwent only handling without exposure to the behavioral paradigm. On day 1, mice went through a habituation session in context A, equipped with an arena with white walls and a stainless-steel foot shock grid. White light was used for illumination (250 lx) and the fan power was adjusted to 30 %. A small amount of 70 % ethanol (v/v) was wiped through the bottom of the arena to generate a weak background odor. In combination with context A, animals were transferred individually in a transport cage filled with bedding material to the experimental room. After placing the animal into the arena, each animal was allowed to habituate for 10 min to context A without any stimulus presentation. The arena was cleaned with soap and water between the individuals during all test sessions. On day 2, fear conditioning was implemented in context A. Therefore, the foot shock grid was connected to a shocker-scrambler unit. After a 2 min baseline period (Bl), animals received five pairings of a 30 s pure tone (conditioned stimulus, CS, 7.5 kHz, 60 dB) co-terminating with a 1 s foot shock (unconditioned stimulus, US, 0.36 mA). The inter-trial intervals (ITIs) between the stimulus presentations ranged between 30–120 s. After the last stimulus was presented, animals remained in the arena for additional 60 s as a post-stimulus time (PST). On day 3, fear retrieval and extinction occurred in context B, equipped with black and white striped walls and a perforated floor plate. Illumination was given with red light (28 lx) and the fan power was set to 100 %. A weak odor was created by wiping the bottom of the arena with a small amount of 1 % acetic acid (v/v). In combination with context B, the transport cage for animal transfer to the experimental room was embedded with paper towels. The stimulus protocol consisted of a 2 min Bl period, followed by 14 CS presentations separated by individual ITIs with comparable length to the conditioning session. The extinction session was completed with an PST of 60 s.

### Behavioral analysis

Video recording and stimulus presentation was controlled with a custom software written in MATLAB (MathWorks). Post hoc video analysis was performed with EthoVision XT tracking software (11.5, Noldus, Wageningen, Netherlands). Freezing behavior was defined as the absence of any movement except respiration with a duration of at least two seconds. Immobility behavior was detected if less than 1 % of the defined animal surface was moving. The percentage of freezing and immobility behavior was calculated for different time intervals (Bl, CS, ITI and PST) and was binned as remarked in each figure. The first CS presentation of the conditioning session was not included. The fear memory retrieval test on day 2 reflects an average of the first two CS presentations. The maximum movement velocity was analyzed as an indicator of foot shock reactivity (38). Due to technical issues, one 2CKO animal was not recorded during the habituation session. This animal was not included in the analysis of distance moved (Fig. 1 h).

**Figure 1.**
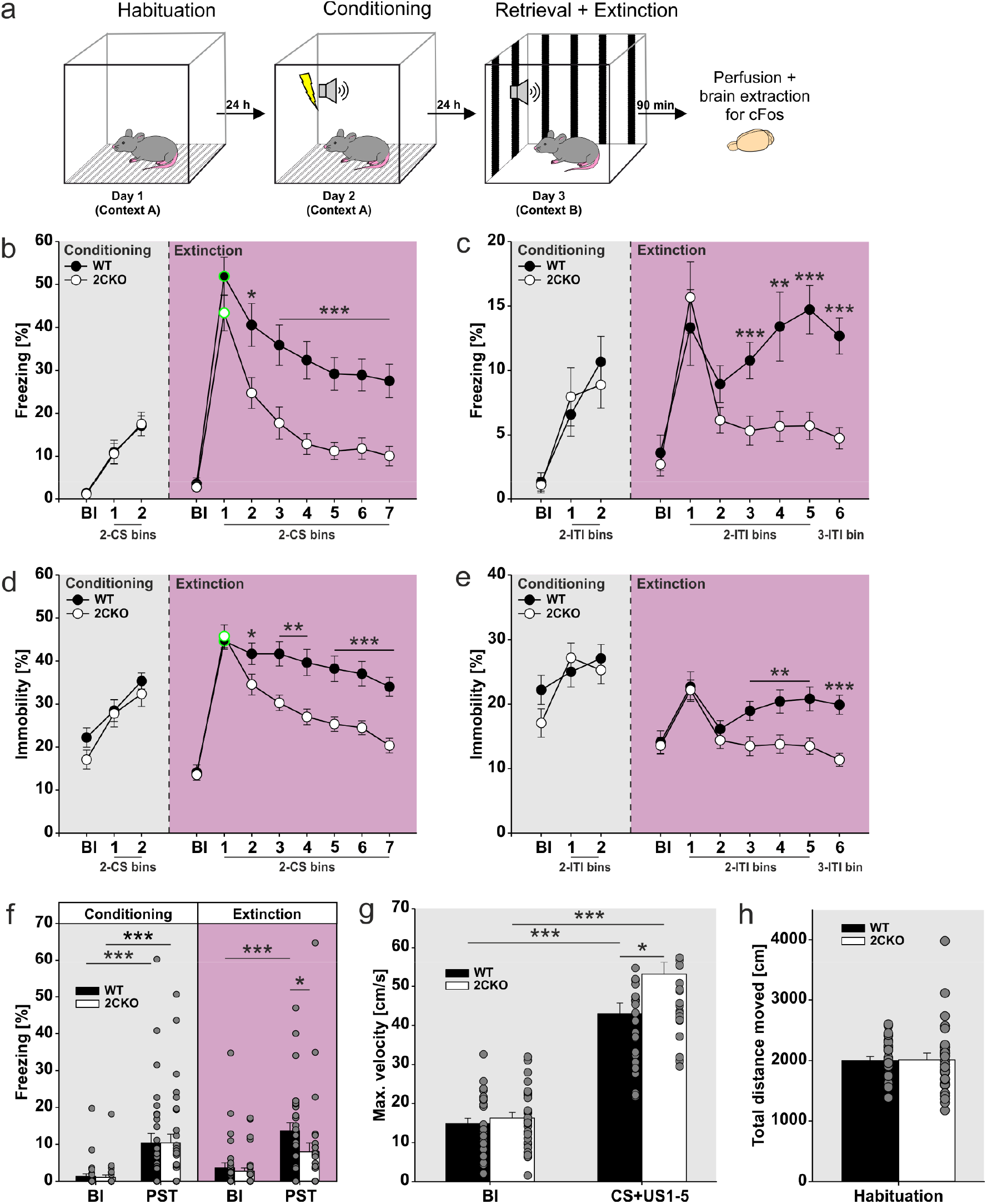
Constitutive 5-HT2CR knock-out facilitates fear extinction. (**a**) Schematic representation of the experimental paradigm. (**b**) Freezing behavior during conditioning (day 2) and extinction (day 3) session in respect to conditioned stimulus (CS) time bins and baseline (Bl) period. Circles outlined in green reflect the fear retrieval test (averaged freezing to the first two CS of the extinction session). 2CKO mice showed less CS-freezing during extinction: Mann-Whitney Rank Sum test (WT vs. 2CKO); bin2(CS): P = 0.023, bin3(CS): P = 0.001, bin4–bin7(CS): P = < 0,001. (**c**) Freezing behavior during conditioning (day 2) and extinction (day 3) session in respect to inter-trial interval (ITI) time bins and baseline (Bl) period. 2CKO mice showed less ITI-freezing during extinction: Mann-Whitney Rank Sum test (WT vs. 2CKO); bin3(ITI): P = 0.001, bin4(ITI): P = 0.009, bin5–6(ITI): P = < 0.001. (**d**) Immobility behavior during conditioning (day 2) and extinction (day 3) session in respect to conditioned stimulus (CS) time bins and baseline (Bl) period. Circles outlined in green reflect the fear retrieval test (averaged immobility to the first two CS of the extinction session). 2CKO mice showed less CS-immobility during extinction: Mann-Whitney Rank Sum test (WT vs. 2CKO); bin2(CS): P = 0.033, bin3(CS): P = 0.002, bin4(CS): P = 0.003, bin5(CS): P = 0.001, bin6–7(CS): P = < 0.001. (**e**) Immobility behavior during conditioning (day 2) and extinction (day 3) session in respect to inter-trial interval (ITI) time bins and baseline (Bl) period. 2CKO mice showed less ITI-immobility during extinction: Mann-Whitney Rank Sum test (WT vs. 2CKO); bin3(ITI): P = 0.003, bin4– 5(ITI): P = 0.004, bin6(ITI): P = < 0.001. (**f**) Comparison of baseline (Bl) and post-stimulus time (PST) freezing during conditioning (day 2) and extinction (day 3) session. Conditioning procedure increased freezing in both genotypes: Wilcoxon Signed Rank test (Bl vs. PST); WT: P = < 0.001, 2CKO: P = < 0.001. WT mice showed higher freezing in extinction PST: Wilcoxon Signed Rank test (Bl vs. PST); WT: P = < 0.001. Mann-Whitney Rank Sum test (WT vs. 2CKO); PST: P = 0.011. (**g**) Maximum movement velocity as unconditioned stimulus (US, foot shock) reactivity. Movement velocity was compared between baseline (BL) and CS+US time, whereby US was presented during the last second of each 30 s CS interval of the conditioning session (day 2).US presentation increased movement velocity in both genotypes: Wilcoxon Signed Rank test (Bl vs. PST); WT: P = < 0.001, 2CKO: P = < 0.001. 2CKO mice showed a higher US reactivity: Mann-Whitney Rank Sum test (WT vs. 2CKO); CS+US1-5: P = 0.021. (**h**) Total distance moved during habituation (day 1) was similar in both genotypes. On freezing and immobility time curves (b–e), each bin reflects two to three averaged time intervals depicted underneath the x-axis. WT mice (n = 29), 2CKO mice (n = 30, for total distance moved in (h) n = 29). Data are shown as means ± SEM. *P < 0.05, **P < 0.01, ***P < 0.001.

### Surgical procedure for virus and tracer injections

For anterograde labeling of neuronal terminals, the adeno-associated virus AAV1.CAG.FLEX.tdTomato.WPRE.bGH (#59462, Addgene, Watertown, MA, USA) was used. The AAV was injected either into the DRC of ePet1-Cre mice for selective tdTomato expression in serotonergic cells, or in the BNSTad of Gad2-Ires-Cre mice for selective expression in GABAergic cells. The retrograde tracer Fluorogold (1 %, v/v, in NaCl, H22845, Thermo Fischer Scientific, Schwerte, Germany) was injected into the BNSTad or medial DRN 10–11 days prior to the behavioral experiments. Mice were initially anesthetized with 5 % isoflurane (v/v) and placed in a stereotactic frame (Stoelting, Wood Dale, IL, USA). To prevent corneal drying during surgery, the eyes were coated with a moisturizing balm. For maintenance of anesthesia, isoflurane levels were adjusted to 1.5–2.5 % (v/v) for the entire procedure. The body temperature was controlled with a heating pad. Virus and tracer delivery was conducted with customized glass pipets (tip diameter 5–10 µm) via pressure injection. Therefore, a small craniotomy was drilled to target the respective brain area with the following stereotactic coordinates, all related to bregma: for DRC with a 24 ° angle from posterior, AP -6.20 mm, DV -3.60 mm; for medial DRN, 24 ° angle from posterior, AP -6.20 mm, DV -3.80 mm; and for BNSTad, AP +0.26 mm, ML +/- 0.75 mm, DV -4.20 mm. Postoperative care included subcutaneous application of Carpofen 2 mg/kg (Rimadyl, Zoetis, Berlin, Germany). AAV injected animals were kept in their home cages for two weeks to enable sufficient virus expression before perfusion without exposure to the behavioral paradigm.

### Histology and immunohistochemistry

Ninety minutes after fear extinction on day 2, mice were deeply anesthetized and transcardially perfused with ice-cold PBS (1x) followed by 4 % paraformaldehyde in PBS (PFA, w/v, pH 7.4, Sigma Aldrich, Taufkirchen, Germany). The brains were post fixed in 4 % PFA over night at 4°C and cryoprotected in 30 % sucrose (w/v, Sigma Aldrich, Taufkirchen, Germany). Following cryoprotection, brains were frozen and cut into 30 µm thick sections with a cryostat (CM3050 S, Leica, Wetzlar, Germany). All stainings described were performed free-floating. Therefore, 24-well plates (STARLAB, Hamburg, Germany) were filled with PBS for collecting the sections. To get rid of excess embedding material, all slices were rinsed with PBS (3x 10 min).

For combined cFos and TPH2 staining in the DRN, sections were blocked in 10 % normal donkey serum (NDS, v/v, Merck Millipore, Darmstadt, Germany) in 0.3 % PBS-Triton X-100 (PBS-T, v/v, Sigma Aldrich, Taufkirchen, Germany) for 1 h at room temperature (RT) to reduce nonspecific antibody binding. Subsequently, sections were incubated with a primary antibody solution (rabbit anti TPH2 (1:500, ab184505, Abcam, Cambridge, UK) and mouse anti cFos (1:500, sc-271243, Santa Cruz Biotechnology, Dallas, TX, USA)) containing 5 % NDS in 0.3 % PBS-T for ∼17 h at 4 °C. Following incubation, DRN sections were rinsed in PBS (3 × 10 min) and incubated with a secondary antibody solution (donkey anti rabbit Cy5 (1:1000, 711-175-152, Jackson Immunoresearch, Ely, UK) and goat anti mouse Alexa 555 (1:500, A21422, Thermo Fischer Scientific, Schwerte, Germany)) containing 5 % NDS in 0.3 % PBS-T for 1.5 h at RT.

For combined Nissl and cFos staining in the amygdala, rinsed sections were permeabilized with 0.1 % PBS-T for 10 min at RT and then incubated with Neurotrace 435/455 nm (1:300, N21479, Thermo Fisher Scientific, Waltham, MA, USA) diluted in PBS for 20 min at RT. After another 10 min incubation with 0.1 % PBS-T, slices were again rinsed with PBS (2x 5 min). This was followed by incubation in 10 % NDS in PBS for 45 min. The blocking solution was aspirated, and the sections were incubated with a primary antibody solution (rabbit anti cFos (1:800, cat. No. 226003, Synaptic Systems, Göttingen, Germany)) containing 3% NDS in PBS for ∼17 h at 4 °C. The slices were rinsed with PBS (3 × 10 min) and incubated with a secondary antibody solution (1:1000, 711-175-152, Jackson Immunoresearch, Ely, UK)) containing 3 % NDS in PBS for 1.5 h at RT.

For combined cFos and PKCδ staining in the BNST, rinsed slices were incubated in 10 mM sodium citrate (pH 6.0) with 0.05 % Tween-20 (v/v, Sigma Aldrich, Taufkirchen, Germany) for 20 min at RT. After rinsing with PBS (2x 5 min), a blocking solution containing 10 % NDS in 0.3 % PBS-T was applied for 1 h at RT. This was followed by an incubation with a primary antibody solution (rabbit anti PKCδ (1:500, ab182126, Abcam, Cambridge, UK) and mouse anti cFos (1:500, sc-271243, Santa Cruz Biotechnology, Dallas, TX, USA)) consisting of 3 % NDS in 0.3 % PBS-T for ∼38 h at 4 °C. The sections were rinsed in PBS (3 × 10 min) and then incubated with a secondary antibody solution (donkey anti rabbit Cy5 (1:1000, 711-175-152, Jackson Immunoresearch, Ely, UK)) and donkey anti mouse Dylight 550 (1:500, SA5-10167, Thermo Fisher Scientific, Waltham, MA, USA)) containing 3 % NDS in 0.3 % PBS-T for 1.5 h at RT.

Following incubation with the respective secondary antibodies, slices from all series were rinsed with PBS (3 × 10 min) and mounted on slides. The slides were coverslipped with ROTI Mount FluorCare medium (Carl Roth, Karlsruhe, Germany) with or without DAPI stain.

### Imaging and analysis

All immunolabeled brain sections were captured with a Leica TCS SP5II confocal laser scanning microscope (Leica Microsystems, Wetzlar, Germany) by using 10x/0.3 NA and 20x/0.7 NA objectives. Sequential Z-stack images with 5–10 optical planes comprising the detectable fluorescence were taken. The images were processed with ImageJ software (39) (1.50i). Cell counting was performed manually with the ImageJ counting tool by two experimenters.

For analysis of the DRN, the typical pattern of tryptophan hydroxylase 2-expressing (TPH2+) cells was used to classify the slices according to three distinct levels (rostral, medial, caudal) on the rostrocaudal AP axis (rostral: -4.42 mm to -4.54 mm from bregma, medial: -4.54 mm to -4.66 mm from bregma, caudal anterior: -4.78 mm to -4.90 mm from bregma, caudal posterior: -4.90 mm to -5.02 mm from bregma). For the caudal level, two portions (caudal anterior and caudal posterior) with distinct subregions were analyzed. A mask depicting the boundaries of individual DRN subregions in the respective level was added to each scanned section. Analyzed subregions include dorsal (DRD), ventral (DRV), ventrolateral (DRVL), interfascicular (DRI) and caudal (DRC) parts, and the ventrolateral periaqueductal gray (VLPAG). Within the same level, the total number of analyzed cells was averaged over 1–3 sections for each animal. Only for the caudal DRN (Fig. 2d), one 2CKO animal of the extinction group was excluded due to tissue damage through histological processing.

**Figure 2.**
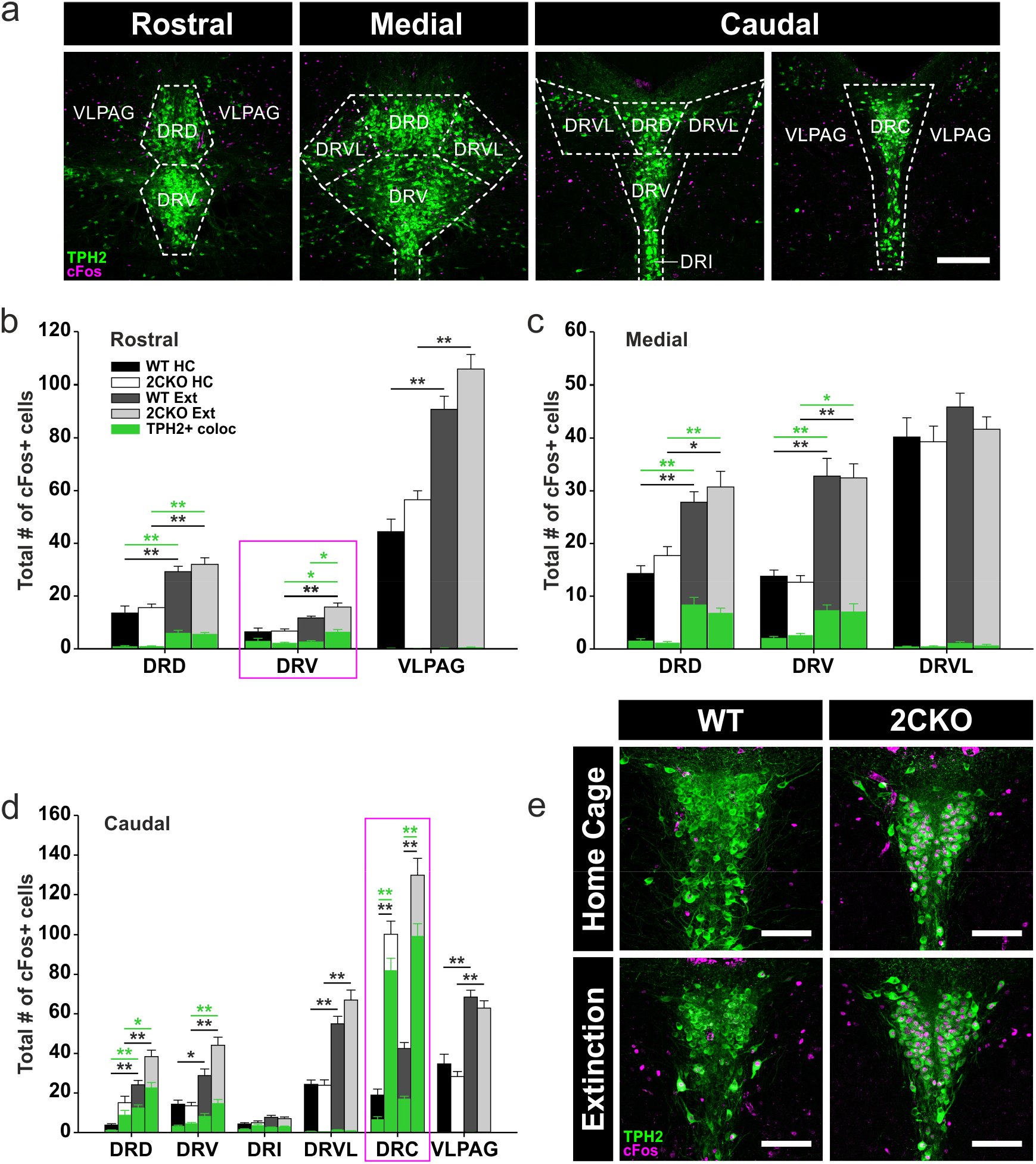
Neuronal activity in the DRN is altered in 2CKO mice. (**a**) Representative images of the DRN levels analyzed. The pattern of TPH2+ 5-HT cells (green) was used to define the respective level. Boundaries are outlined by dashed lines. DRD = dorsal raphe nucleus, dorsal part; DRV = dorsal raphe nucleus, ventral part; DRI = dorsal raphe nucleus, interfascicular part; DRC = dorsal raphe nucleus, caudal part; DRVL = dorsal raphe nucleus, ventrolateral part; VLPAG = ventrolateral periaqueductal gray. Scale bar = 250 µm. (**b**) cFos quantification in the rostral DRN. Significant effects DRD: Kruskal-Wallis one-way ANOVA on ranks (cFos+): P = < 0.001; pairwise Dunn’s test: WT HC vs. WT Ext: P = < 0.01, 2CKO HC vs. 2CKO Ext: P = < 0.01. Kruskal-Wallis one-way ANOVA on ranks (TPH2+/cFos+): P = < 0.001; pairwise Dunn’s test: WT HC vs. WT Ext: P = < 0.01, 2CKO HC vs. 2CKO Ext: P = < 0.01. Significant effects DRV: Kruskal-Wallis one-way ANOVA on ranks (cFos+): P = < 0.001; pairwise Dunn’s test: 2CKO HC vs. 2CKO Ext: P = < 0.01. Kruskal-Wallis one-way ANOVA on ranks (TPH2+/cFos+): P = 0.004; pairwise Dunn’s test: WT HC vs. WT Ext: P = < 0.05, WT Ext vs. 2CKO Ext: P = < 0.05. Significant effects VLPAG: Kruskal-Wallis one-way ANOVA on ranks (cFos+): P = < 0.001; pairwise Dunn’s test: WT HC vs. WT Ext: P = < 0.01, 2CKO HC vs. 2CKO Ext: P = < 0.01. (**c**) cFos quantification in the medial DRN. Significant effects DRD: Kruskal-Wallis one-way ANOVA on ranks (cFos+): P = < 0.001; pairwise Dunn’s test: WT HC vs. WT Ext: P = < 0.01, 2CKO HC vs. 2CKO Ext: P = < 0.05. Kruskal-Wallis one-way ANOVA on ranks (TPH2+/cFos+): P = < 0.001; pairwise Dunn’s test: WT HC vs. WT Ext: P = < 0.01, 2CKO HC vs. 2CKO Ext: P = < 0.01. Significant effects DRV: Kruskal-Wallis one-way ANOVA on ranks (cFos+): P = < 0.001; pairwise Dunn’s test: WT HC vs. WT Ext: P = < 0.01, 2CKO HC vs. 2CKO Ext: P = < 0.01. Kruskal-Wallis one-way ANOVA on ranks (TPH2+/cFos+): P = < 0.001; pairwise Dunn’s test: WT HC vs. WT Ext: P = < 0.01, 2CKO HC vs. 2CKO Ext: P = < 0.05. (**d**) cFos quantification in the caudal DRN. Significant effects DRD: Kruskal-Wallis one-way ANOVA on ranks (cFos+): P = < 0.001; pairwise Dunn’s test: WT HC vs. WT Ext: P = < 0.01, 2CKO HC vs. 2CKO Ext: P = < 0.01. Kruskal-Wallis one-way ANOVA on ranks (TPH2+/cFos+): P = < 0.001; pairwise Dunn’s test: WT HC vs. WT Ext: P = < 0.01, 2CKO HC vs. 2CKO Ext: P = < 0.05. Significant effects DRV: Kruskal-Wallis one-way ANOVA on ranks (cFos+): P = < 0.001; pairwise Dunn’s test: WT HC vs. WT Ext: P = < 0.05, 2CKO HC vs. 2CKO Ext: P = < 0.01. Kruskal-Wallis one-way ANOVA on ranks (TPH2+/cFos+): P = < 0.001; pairwise Dunn’s test: 2CKO HC vs. 2CKO Ext: P = < 0.01. Significant effects DRVL: Kruskal-Wallis one-way ANOVA on ranks (cFos+): P = < 0.001; pairwise Dunn’s test: WT HC vs. WT Ext: P = < 0.01, 2CKO HC vs. 2CKO Ext: P = < 0.01. Significant effects DRC: Kruskal-Wallis one-way ANOVA on ranks (cFos+): P = < 0.001; pairwise Dunn’s test: WT HC vs. 2CKO HC: P = < 0.01, WT Ext vs. 2CKO Ext: P = < 0.01. Kruskal-Wallis one-way ANOVA on ranks (TPH2+/cFos+): P = < 0.001; pairwise Dunn’s test: WT HC vs. 2CKO HC: P = < 0.01, WT Ext vs. 2CKO Ext: P = < 0.01. Significant effects VLPAG: Kruskal-Wallis one-way ANOVA on ranks (cFos+): P = < 0.001; pairwise Dunn’s test: WT HC vs. WT Ext: P = < 0.01, 2CKO HC vs. 2CKO Ext: P = < 0.01. (**e**) Representative immuno-stained DRC sections. cFos+ cells (magenta) showed a high colocalization with TPH2+ 5-HT cells (green) in 2CKO animals under home cage and extinction conditions. Scale bars = 100 µm. Boxes in magenta highlight subregions with significant genotype effects. HC: WT mice (n =10), 2CKO mice (n = 11); Ext: WT mice (n = 13), 2CKO mice (n = 13, for caudal in (d) n = 12). Data are shown as means ± SEM. *P < 0.05, **P < 0.01, ***P < 0.001.

For analysis of the amygdala, boundaries of the subregions were outlined manually using Nissl staining in combination with the mouse brain atlas (40). Three separate levels (rostral, medial, caudal) on the rostrocaudal AP axis, each encompassing 300 µm, were analyzed (rostral: -0.90 mm to -1.20 mm from bregma, medial: -1.50 mm to -1.80 mm from bregma, caudal: -1.90 mm to -2.20 mm from bregma). The amygdala nuclei examined included the basolateral amygdala (BLA) with the subregions basal amygdala (BA), posterior portion of the basolateral amygdala (BLp) and lateral amygdala (LA), and the central amygdala nucleus (CeA) containing a lateral (CeL) and a medial (CeM) portion. The analysis of the CeA was only restricted to the medial level. Within each level, the total number of counted cells was averaged over 3–7 sections from both hemispheres for each animal.

Analysis of the BNST was restricted to two subregions of the dorsal BNST, the oval nucleus (BNSTov) and the anteriodorsal subregion (BNSTad). The protein kinase C δ (PKCδ) staining was used as a marker for the BNSTov (41, 42). The BNSTad was defined by anatomical boundaries including the anterior commissure (ventral), the juxta capsular nucleus and the internal capsule (lateral) and the lateral septum (medial). For each animal, the total number of cells was counted in 6–12 sections (between +0.10 mm and +0.30 mm from bregma) from both hemispheres and averaged.

### Statistics

Statistical analysis was performed with SigmaPlot 12.5 (Systat Software). All data was tested for normal distribution via Shapiro-Wilk test and equal variances were assessed via Levene’s test. As most of the data was not normally distributed or equal variance test failed, non-parametric tests were mainly used to reveal significances. For behavioral analysis (Fig. 1b-e), between group effects were analyzed pairwise by using the Mann-Whitney Rank Sum test. Wilcoxon Signed Rank test was used to reveal within groups differences at distinct time points (Fig. 1f and 1g). Analysis of cFos in the DRN and the amygdala was performed via Kruskal-Wallis one-way ANOVA on ranks followed by pairwise multiple comparison procedure using Dunn’s method. Parametric analysis was used for cFos evaluation in the BNST (Fig. 3b) via two-way ANOVA and Holm-Sidak post hoc comparison. Further parametric student’s t-test (two-tailed) was used for combined cFos and FG tracer analysis in the DRC (Fig. 5e) to reveal between group effects. Final animal numbers are mentioned in the figure legends. For all results, the level of significance was set to P < 0.05. All data are expressed as mean ± SEM.

**Figure 3.**
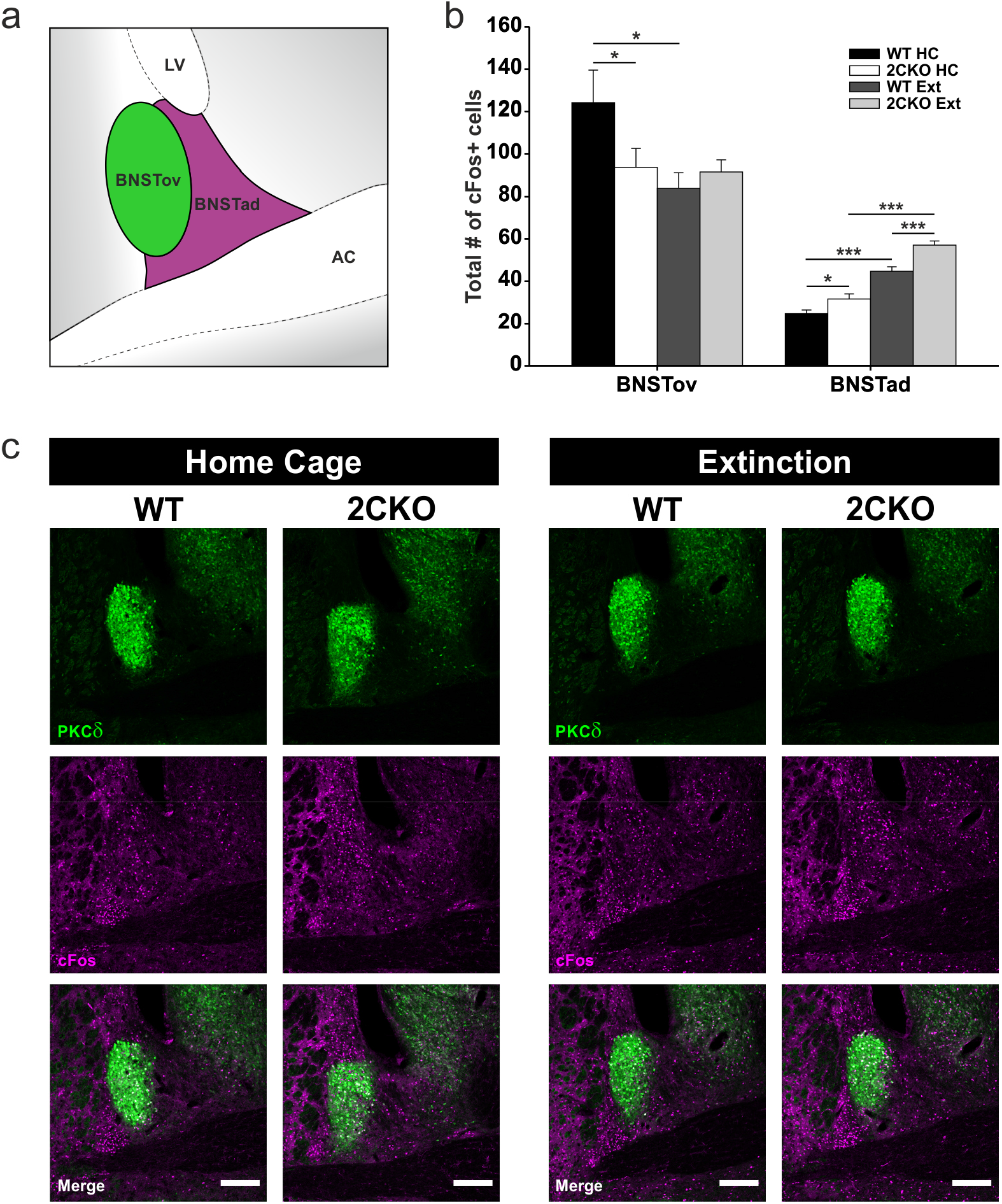
Altered neuronal activity in the dorsal BNST supports faster fear extinction in 2CKO mice. (**a**) Schematic illustration of the dorsal BNST subregions analyzed. BNSTov = bed nucleus of the stria terminalis, oval nucleus; BNSTad = bed nucleus of the stria terminalis, anteriodorsal part; LV = lateral ventricle; AC = anterior commissure. (**b**) cFos quantification in the BNSTov and BNSTad. In the BNSTov, 2CKO mice showed reduced cFos levels under home cage conditions and extinction treatment reduced cFos in WT mice: Two-way ANOVA (treatment): F_(1,19)_ = 4.973, P = 0.038; pairwise Holm-Sidak test: WT HC vs. 2CKO HC: P = 0.036, WT HC vs. WT Ext: P = 0.011. In the BNSTad, 2CKO mice showed increased cFos levels under home cage and extinction conditions and extinction treatment increased cFos in both genotypes: Two-way ANOVA (genotype): F_(1,19)_ = 20.736, P = < 0.001; Two-way ANOVA (treatment): F_(1,19)_ = 114.923, P = < 0.001; pairwise Holm-Sidak test: WT HC vs. 2CKO HC: P = 0.034, WT Ext vs. 2CKO Ext: P = < 0.001, WT HC vs. WT Ext: P = < 0.001, KO HC vs. KO Ext: P = < 0.001. (**c**) Representative immuno-stained BNST sections. PKCδ (green), used as a marker for the BNSTov subregion, in combination with cFos (magenta). Scale bars = 200 µm. HC: WT mice (n = 5), 2CKO mice (n = 6); Ext: WT mice (n = 5), 2CKO mice (n = 7). Data are shown as means ± SEM. *P < 0.05, **P < 0.01, ***P < 0.001.

## Results

### Constitutive knock-out of the 5-HT2CR facilitates fear extinction

To determine the behavioral consequences of 5-HT2CR knock-out, 2CKO and WT mice were exposed to an auditory fear conditioning and extinction protocol (Fig. 1a). Freezing behavior was analyzed for CS (Fig. 1b) or ITI (Fig. 1c) time bins during the conditioning (day 2) and extinction (day 3) session. Statistical analysis of the data revealed no significant alterations in freezing behavior during fear conditioning between WT and 2CKO groups. Freezing during fear memory retrieval test (Fig. 1b, bin1) was also not significantly different. However, freezing during the extinction session was significantly decreased in 2CKO mice (Fig. 1b and 1c), reflecting a faster fear extinction in 2CKO animals. Besides freezing, we analyzed the immobility behavior throughout the fear conditioning paradigm (Fig. 1d and 1e). For fear conditioning, statistical analysis indicated no significant differences in immobility between WT and 2CKO groups. Similarly, immobility behavior during fear memory retrieval test (Fig. 1d, bin1) was not significantly altered in 2CKO animals. In contrast, 2CKO mice showed significantly less immobility behavior during the extinction session in comparison to WT littermates (Fig. 1d and 1e), supporting the effect of faster fear extinction in 2CKO animals. Comparison of baseline (BL) freezing to post-stimulus time (PST) freezing (Fig. 1f) revealed significant within group effects through the conditioning session due to successful fear acquisition. PST freezing during the extinction session was significantly reduced in 2CKO animals when compared to WT controls (Fig. 1f). Freezing level at PST was not significantly different from Bl freezing in 2CKO mice, indicating a successful fear extinction. For WT animals, freezing was still significantly increased at PST when compared to Bl (Fig. 1f), suggesting that fear memory in WT animals was not fully extinguished at the end of the extinction session. Further, we examined the animal’s reactivity to the unconditioned foot shock (US), by analyzing the maximum movement velocity (Fig. 1g) (38). The US presentation at the end of the CS interval increased the maximum movement velocity significantly within both groups, when compared to Bl period (Fig. 1g). Between group analysis indicated a significant higher velocity of 2CKO mice due to US presentation when compared to WT littermates. These results are in line with the findings of Bonasera et al. (43), describing increased affective responses to unconditioned stimuli in 2CKO animals, but notably, the pain sensitivity to noxious stimuli was not significantly altered in 2CKO mice. Moreover, statistical analysis of the total distance moved during the habituation session (Fig. 1h) did not reveal significant differences. This indicates that 2CKO animals did not display hyperactive behavior, at least in our paradigm. Taken together, our behavioral analysis revealed that the constitutive knock-out of the 5-HT2CR facilitates fear extinction in an auditory fear conditioning paradigm.

### Basal activity of 5-HT neurons is increased in the DRC subregion of the DRN in 2CKO mice

To assess if faster fear extinction in 2CKO mice is associated with changes in neuronal activity due to 5-HT2CR knock-out, we mapped cFos expression in different brain areas. Thus, 90 min after fear extinction (day 3), mice from WT (WT Ext) and 2CKO (2CKO Ext) groups were perfused (Fig. 1a). Additionally, WT (WT HC) and 2CKO (2CKO HC) mice that permanently remained in their home cages were perfused as controls. After extraction of the brains, the tissue was immunohistochemically processed for cFos. In the first step, cFos expression was quantified in the DRN, the main source of serotonergic forebrain innervation (44). The DRN accounts for the largest number of serotonergic neurons and is generally considered to play an important role in the processing of anxiety and fear (1, 45, 46). As 5-HT neurons are topographically organized in the DRN (36), our analysis included three distinct levels on the rostrocaudal extend (rostral, medial, caudal), each encompassing several subregions (see Fig. 2a). Only the caudal DRN level contained two distinct portions. Besides the exclusive analysis of cFos positive (cFos+) cells, we analyzed cFos colocalization with tryptophan hydroxylase 2 positive (TPH2+) cells, to mark neuronal activity in 5-HT neurons. The statistical evaluation indicated significant treatment effects (home cage vs. extinction) within the 2CKO and WT groups in various DRN subregions (Fig 2b, 2c and 2d). In contrast, significant genotype effects (WT vs. 2CKO) could be detected exclusively in two subregions, in the DRV of the rostral DRN (Fig. 2b, magenta box), the most significant genotype effect was found in the DRC of the caudal DRN (Fig. 2d, magenta box). For the rostral DRV, extinction treatment had no effect on cFos in WT mice, whereas 2CKO mice displayed increased cFos levels in TPH2+ 5-HT cells following extinction. Similarly, 2CKO animals displayed elevated cFos levels in the caudal DRC when compared to WT littermates. Interestingly, this genotype effect was independent of extinction treatment. Moreover, the vast majority of cFos+ cells in the DRC of 2CKO mice showed a colocalization with TPH2 (Fig. 2d and 2e). Taken together, our cFos analysis in the DRN revealed increased 5-HT cellular activity in the rostral DRV and the caudal DRC in 2CKO mice. Enhanced neuronal activity in the DRV was extinction-associated, whereas neuronal activity in the DRC was altered under basal conditions. Thus, we assume that 2CKO mice exhibit an increased 5-HT release in DRC target areas in general.

### Neuronal activity in the amygdala is not altered in 2CKO mice

In a second step, we quantified cFos expression in the amygdala due to its close interaction with the serotonergic system and its major position in the fear circuitry. Our amygdala analysis included three distinct levels (rostral, medial, caudal) on the rostrocaudal extend (see Supplementary Fig. 1a). The statistical analysis revealed significant treatment effects (home cage vs. extinction) within the 2CKO and WT groups in several amygdala subregions (Supplementary Fig. 1b–1e). Surprisingly, no significant genotype effects (WT vs. 2CKO) could be detected in any of the subregions analyzed. To conclude, extinction-associated cFos evaluation in the amygdala indicated no alterations in neuronal activity in 2CKO mice in comparison to WT mice.

### Faster fear extinction in 2CKO mice is associated with altered neuronal activity in the dorsal BNST

After our cFos results revealed no alterations in neuronal activity in the amygdala in 2CKO mice, we focused on the BNST as a part of the extended amygdala. Our analysis was restricted to the dorsal BNST and included two distinct subregions, the BNSTov and the BNSTad (see Fig. 3a). An immunohistochemical staining against protein kinase C δ (PKCδ) was used to define the BNSTov subdivision (shown in Fig. 3c). The BNSTad was defined by anatomical landmarks. For BNSTov, statistical analysis revealed a significant genotype effect (WT vs. 2CKO) exclusively for the HC condition (Fig. 3b). Here, cFos levels were significantly decreased in 2CKO mice, indicating reduced neuronal activity under basal conditions. Moreover, we found an extinction-induced significant reduction in cFos expression for WT mice in the BNSTov, whereas extinction treatment had no effect on 2CKO animals. For BNSTad, our analysis revealed significant genotype differences for home cage and extinction conditions (Fig. 3b). 2CKO animals displayed increased cFos levels in both treatment conditions when compared to WT animals. Further, 2CKO and WT mice showed an extinction-associated elevation of cFos in the BNSTad. These results suggest that extinction in WT animals is associated with two opposite effects in the dorsal BNST. Firstly, a decrease of neuronal activity in BNSTov and secondly an increase of activity in BNSTad. In conclusion, neuronal activity is altered in 2CKO animals in an extinction-supporting direction, even under basal home cage conditions.

### The DRN and the dorsal BNST are reciprocally connected

The cFos evaluation indicated altered neuronal activity in the DRN and the BNST of 2CKO mice. Hence, we next investigated the DRN-BNST connectivity. As we found changes in basal activity in both structures, we decided to focus on the DRC subregion of the DRN. For anterograde labeling of neuronal terminals, a viral approach based on an AAV (AAV1.CAG.FLEX.tTomato.WPRE.bGH) carrying a double-floxed tdTomato transgene was used. In the first step, the AAV was injected into the DRC of an ePet1-Cre mouse, to label 5-HT terminals innervating the BNST (Fig. 4a). Verification of tdTomato positive (tdTomato+) cells in the DRN revealed that the expression was selectively restricted to TPH2+ 5-HT cells in the caudal DRN level (Fig. 4b and Supplementary Fig. 2). Indeed, our results revealed a serotonergic innervation of the BNSTad (Fig. 4c and 4d). The majority of tdTomato+ 5-HT terminals was localized in the BNSTad, whereas only a few terminals were found in the BNSTov. Since GABAergic neurons represent the largest population in the BNST (47), the AAV was injected into the BNSTad of a Gad2-Ires-Cre mouse in a second step, to verify if the BNSTad and the DRN are reciprocally connected (Fig. 4a and 4e). Again, the DRN was analyzed at distinct levels due to its topographical organization (Fig. 4f). Our results indicated an overall innervation of the DRN by GABAergic terminals arising from the BNSTad. The majority of tdTomato-expressing GABAergic fibers terminated in the medial DRN, whereas the caudal levels including the DRC were only sparsely innervated. In conclusion, our AAV-based labeling of anterograde terminals revealed a serotonergic DRC-BNSTad projection. GABAergic neurons in the BNSTad in turn project to multiple levels of the DRN.

**Figure 4.**
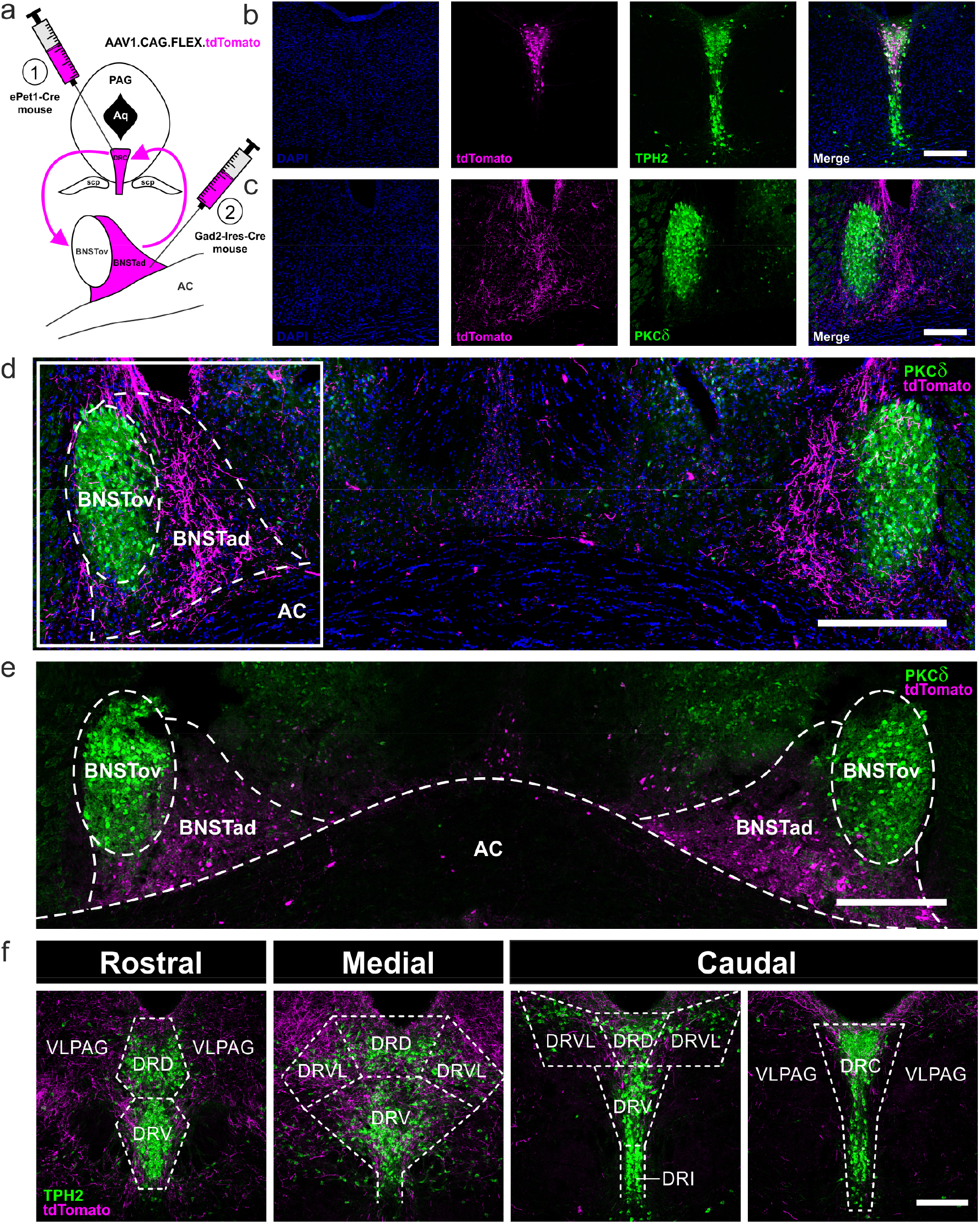
The DRN and the dorsal BNST are reciprocally connected. (**a**) AAV-based labeling strategy of anterograde terminals. AAV1.CAG.FLEX.tdTomato virus was injected either into the DRC of an ePet1-Cre mouse (1) to label 5-HT terminals innervating the BNST, or into the BNSTad subregion of a Gad2-Ires-Cre mouse (2) to label GABAergic terminals innervating the DRN. (**b**) DRC injection site in an ePet1-Cre mouse indicates selective tdTomato expression (magenta) in TPH2+ 5-HT cells (green). Scale bar = 200 µm. (**c**) BNST target site in the ePet1-Cre mouse indicates that tdTomato+ 5-HT terminals (magenta) innervate primarily the BNSTad subregion. PKCδ (green) was used as a marker for the BNSTov subregion. Scale bar = 200 µm. (**d**) Overview of the dorsal BNST in the ePet1-Cre mouse reflecting the bilateral innervation of the BNSTad with tdTomato+ 5-HT terminals (magenta) arising from the DRC. PKCδ (green) was used as a marker for the BNSTov subregion. Boxed region is separately shown in (c). Scale bar = 300 µm. (**e**) Overview of bilateral BNSTad injection sites in a Gad2-Ires-Cre mouse. tdTomato+ GABAergic cells (magenta). PKCδ (green) was used as a marker for the BNSTov subregion. Scale bar 300 µm. (**f**) DRN target site in the Gad2-Ires-Cre mouse indicates that tdTomato+ GABAergic terminals (magenta) arising from the BNSTad innervate all rostrocaudal DRN levels. Strongest innervation was found on the medial level. The pattern of TPH2+ 5-HT cells (green) was used to define the respective level. Boundaries are outlined by dashed lines. DRD = dorsal raphe nucleus, dorsal part; DRV = dorsal raphe nucleus, ventral part; DRI = dorsal raphe nucleus, interfascicular part; DRC = dorsal raphe nucleus, caudal part; DRVL = dorsal raphe nucleus, ventrolateral part; VLPAG = ventrolateral periaqueductal gray. Scale bar = 250 µm.

### A feedback projection from the BNSTad to the DRC does not contribute to faster fear extinction in 2CKO mice

Finally, we used the retrograde tracer Fluorogold (FG) in combination with cFos mapping to examine if interactions between the DRN and the BNST could contribute to faster fear extinction in 2CKO mice. Therefore, 2CKO and WT mice were injected with 1 % FG either into the BNSTad or into the medial DRN (Fig 5a). Since our viral tdTomato approach revealed the medial DRN as the main termination site for GABAergic BNSTad projection neurons, FG injections were targeted to this DRN level. Additionally, our preliminary data (unpublished) indicated that FG injections into the DRC fail to label cells in BNSTad. Ten to eleven days after FG was injected, the animals underwent the fear conditioning and extinction procedure (Fig. 5b). Perfusion took place 90 min after extinction exposure. The brains were extracted and immunohistochemically processed for cFos. The FG injection into the BNSTad resulted in dense retrograde labeling of FG positive (FG+) 5-HT cells (TPH2+) in the caudal DRC (Fig 5c). Only few FG+ 5-HT cells were found at other DRN levels through the rostrocaudal extend (Supplementary Fig. 3). Thus, our retrograde labeling confirmed the discrete serotonergic DRC-BNSTad projection. Moreover, cFos analysis in combination with TPH2 staining revealed a highly significant genotype effect (Fig. 5e). The number of co-labeled (FG+/cFos+) and triple-labeled (FG+/cFos+/TPH2+) cells was increased in 2CKO mice when compared to WT littermates. These results are in line with our initial cFos study and confirm that a discrete serotonergic DRC-BNSTad projection exhibits enhanced activity in 2CKO mice. The FG injection in the medial DRN resulted in retrograde labeling of cells primarily located in the BNSTad subregion (Fig. 5d). Only a small number of FG+ cells was localized in the BNSTov (Fig. 5d and 5f). Noteworthy, the combined analysis with cFos (FG+/cFos+) revealed no significant genotype differences, indicating that BNSTad-DRN projection neurons do not display extinction-associated activity changes. Taken together, our retrograde tracer approach confirmed enhanced activity of a discrete serotonergic DRC-BNSTad pathway in 2CKO mice. Moreover, the extinction-induced activity increase in the BNSTad subregion is not associated with a BNSTad-DRN feedback projection. Thus, faster fear extinction in 2CKO mice may depend on BNSTad projections to other extinction-related downstream areas.

**Figure 5.**
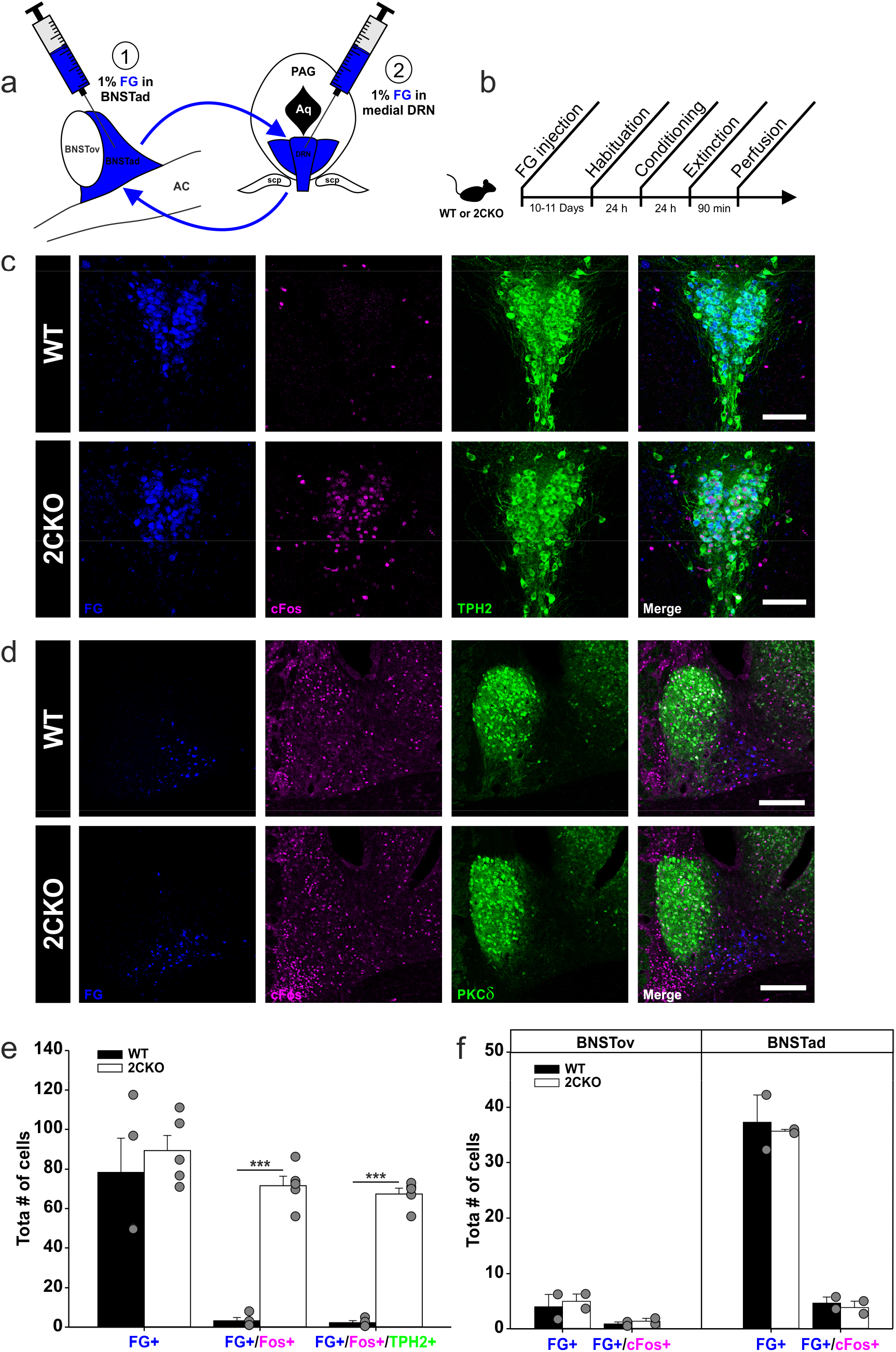
A BNSTad-DRN pathway is not involved in faster fear extinction in 2CKO mice. (**a**) Fluorogold (FG)-based retrograde labeling strategy in 2CKO and WT mice. FG was injected either into the BNSTad (1) for retrograde labeling of DRN cells, or into the medial DRN (2), for retrograde labeling of BNSTad cells. (**b**) Experimental workflow. (**c**) Representative immuno-stained DRC sections of mice injected into the BNSTad. Retrogradely labeled FG cells (blue) are highly colocalized with cFos (magenta) and TPH2 (green) in the 2CKO mouse. Scale bar = 100 µm. (**d**) Representative immuno-stained dorsal BNST sections of mice injected into the medial DRN. Extinction-induced cFos (magenta) is not associated with FG (blue) labeled neurons projecting to the DRN. PKCδ (green) was used as a marker for the BNSTov subregion. Scale bar = 200 µm. (**e**) Evaluation of retrogradely labeled FG cells in the DRN revealed a higher number of FG+/cFos+ and (FG+/cFos+/TPH2+) in 2CKO mice. Two-tailed student’s t-test (WT vs. 2CKO); FG+/cFos+: P = < 0.001, FG+/cFos+/TPH2+: P = < 0.001. WT mice (n = 4), 2CKO mice (n = 5). (**f**) Evaluation of retrogradely labeled FG cells in the dorsal BNST revealed no genotype differences. WT mice (n = 2), 2CKO mice (n = 2). Data are shown as means ± SEM. *P < 0.05, **P < 0.01, ***P < 0.001.

## Discussion

Collectively, our results revealed an enhanced fear extinction effect in 2CKO mice in an auditory fear conditioning paradigm. 2CKO mice displayed altered cFos levels in the DRN and the dorsal BNST, indicating changes in neuronal activity. These alterations were partially extinction-associated, but cFos levels were also changed under basal conditions in 2CKO mice. The assessment of DRN-BNST connectivity identified a serotonergic DRC-BNSTad pathway, showing increased activity in 2CKO animals even under basal conditions. Neuronal activity in the dorsal BNST was shifted in an extinction-supporting direction in 2CKO mice. However, these activity alterations were not associated with a BNSTad-DRN feedback projection.

Previous studies investigating the involvement of 5-HT2CRs in fear and anxiety by using 2CKO mice reported mixed results (48, 49). Our findings are in line with Tecott et al. (50), indicating no alterations in fear retrieval to a tone. A more recent study confirmed the ability of fear retrieval in 2CKO mice by using a context conditioning paradigm (51). Comparable to our data, Nebuka and colleagues showed a faster context extinction in 2CKO mice. In contrast, differences in fear acquisition could not be detected in our paradigm. The researchers postulated that the faster fear extinction may be biased by increased locomotor activity in 2CKO animals. We did neither detect activity changes during the habituation session (day 1), nor during the baseline periods (day 2 and day 3) (Fig. 1h and Supplementary Fig. 4). Thus, we conclude that the fast decline in freezing behavior in 2CKO mice can be estimated as fear reduction, at least in our paradigm. Moreover, Règue and colleagues (52) provided substantial evidence for the involvement of 5-HT2CRs in learned fear responses by using VGV mice. The transgenic VGV strain exhibits an overexpression of the fully edited VGV 5-HT2CR isoform on the cellular surface, causing a phenotype with PTSD-like symptoms. VGV mice display exaggerated fear expression, extensive fear extinction deficits, and fear generalization. This strengthens our finding that a global lack of 5-HT2CRs induces an opposite effect of accelerated fear extinction.

Increased activity of a discrete serotonergic DRC-BNSTad pathway in 2CKO mice reflects a key finding in our study. Within the DRN, 5-HT2CRs are localized on GABAergic interneurons (53, 54) and are described to be involved in negative feedback on 5-HT neurons to regulate anxiety and stress responses (55, 56). Although early studies reported that 5-HT2CR ligands fail to modulate basal 5-HT release in the frontal cortex (57, 58), note that our effect is restricted to a small group of 5-HT neurons, which may not have been considered in the previous studies. Potentially in the caudal DRN, 5-HT2CR isoforms with a high constitutive activity (2, 59) are localized on GABAergic interneurons. If so, the 5-HT2CR knock-out may lead to a disinhibition of serotonergic DRC neurons, reflected by increased activity reported here. Further studies on 5-HT2CR activity in the DRN with respect to the topographical organization of 5-HT neurons will be needed to clarify this assumption.

Our results also uncovered opposite roles of the BNSTov and the BNSTad in fear extinction, as WT mice showed an extinction-associated reduction of activity in the BNSTov and an enhancement in the BNSTad. Optogenetic manipulations revealed similar results in anxiety tasks, indicating an anxiogenic function of the BNSTov, whereas the BNSTad was described to be anxiolytic (15). Matching to this, we found shifted activity in the dorsal BNST in 2CKO mice in an anxiolytic and extinction-supporting direction, even under basal conditions. 2CKO mice have already been described with an anxiolytic phenotype, showing blunted cFos expression in CRF-expressing neurons to anxiety-inducing stimuli (48). As the majority of CRF-expressing neurons is localized in the BNSTov (42), our results support these findings.

Distinct studies, primarily based on SSRIs, revealed a BNST involvement in aversive information processing especially when high 5-HT levels are present (60, 61). It is assumed that under normal physiological conditions, increased 5-HT release in the BNST primarily acts on 5-HT2CRs (56, 62). 5-HT2CR activation in turn engages a BNST microcircuit that inhibits anxiolytic projections to the ventral tegmental area and lateral hypothalamus to increase anxiety (17). In contrast, anxiolytic 5-HT actions in the BNST are attributed to 5-HT2CR desensitization (6) and enhanced 5-HT1AR activation, favoring anxiolytic outputs (63, 64). Additionally, plasticity changes in 5-HT1AR signaling following contextual fear conditioning is thought to buffer the animal’s system against excessive activation (63). Thus, it is likely that in 2CKO mice, an increased 5-HT release into the BNSTad mainly acts on 5-HT1A receptors, supporting the faster extinction effect reported here. Since our tracer results have shown that a reciprocal interaction between the BNSTad and the DRC is not involved in the extinction-facilitating effect in 2CKO mice, further studies identifying BNSTad projections to extinction-associated target sites are needed.

In summary, our data reveal that a global knock-out of 5-HT2CRs is associated with increased activity of a serotonergic DRC-BNSTad pathway, supporting accelerated fear extinction. To the best of our knowledge, this is the first report describing a precise serotonergic DRC-BNSTad pathway, which is altered in a 5-HT2CR-sensitive mechanism. Since anxiolytic effects following systemic long-term SSRI treatment are associated with 2CR desensitization, the effect we reported here may be a key mechanism in SSRI-induced anxiolysis. Finally, fear extinction reflects the core mechanism of exposure-based therapy (EBT) in the treatment of PTSD in humans (65) As EBT is long-lasting and relapse rates are high, augmentation strategies are urgently needed. Thus, our results revealed the 5-HT2CR as a putative drug target for the pharmacological intervention in EBT. Further research will be needed to strengthen this hypothesis.

## Acknowledgements

We would like to thank Stefan Dobers, Winfried Junke, Margaretha Möllmann, Stephanie Krämer, Petra Knipschild, Nicole Ozdovski, Gina Weber, and Manuela Schmidt for excellent technical assistance. We thank Prof. Petra Wahle for her advice regarding histology and IHC techniques. This work was supported by Deutsche Forschungsgemeinschaft (DFG, German Research Foundation) SFB1280 (project number 316803389, project A07) (to K.S. and S.H.), SFB874 (project number 122679504, project B10) (to S.H.), DFG2471/23-1 (to S.H.), DFG2471/21-1 (to S.H.).

## Author contributions

S.T.S., S.H. and K.S. designed the experiments. S.T.S. performed all experiments and analyzed the data. L.M.O. supported the anterograde tracing experiment. D.K. supported cell counting in the amygdala. S.T.S., S.H. and K.S. wrote the manuscript.

---

**Supplementary Figure 1.**
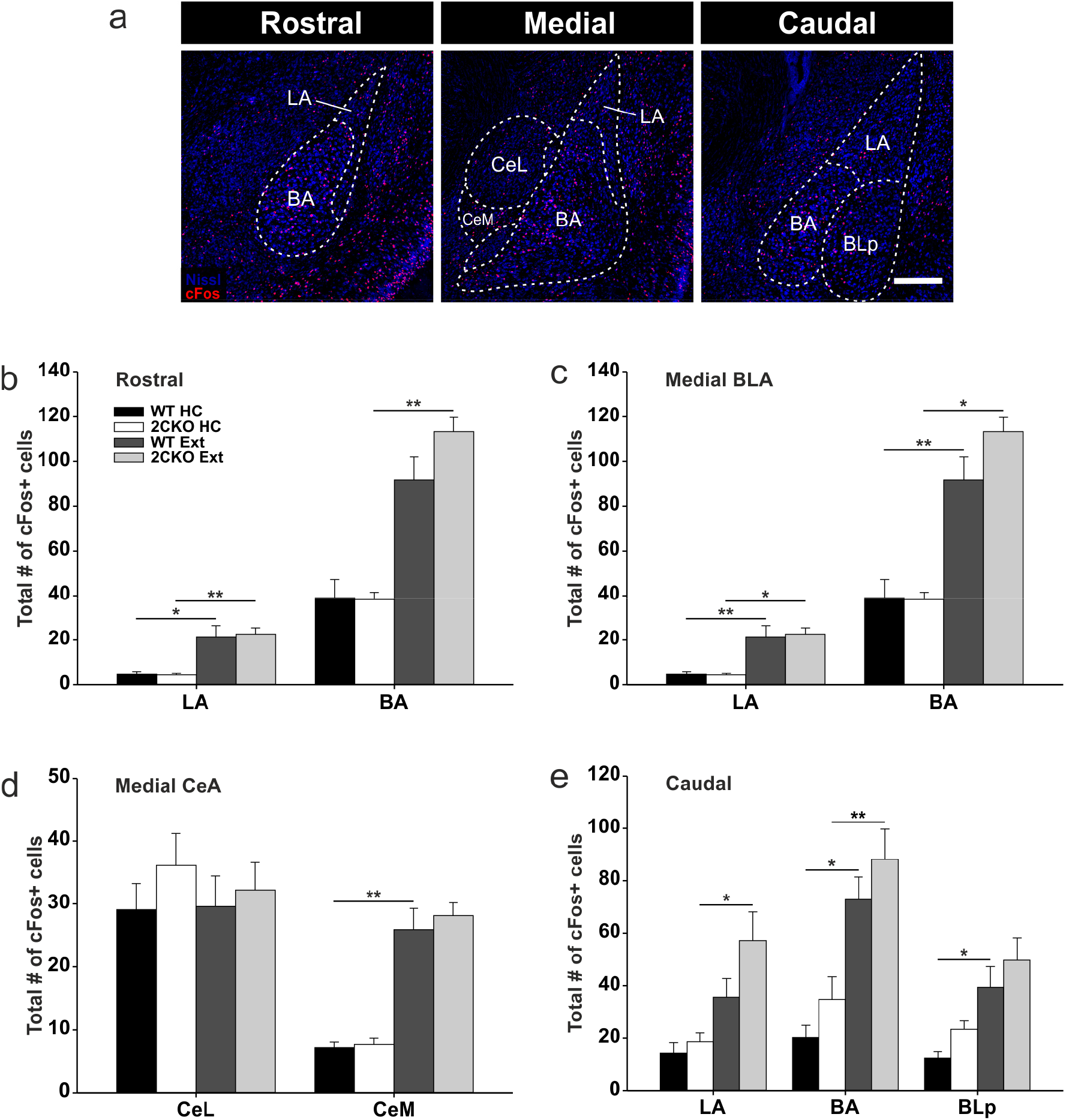
Neuronal activity in the amygdala is not altered in 2CKO mice. (**a**) Representative images of amygdala levels analyzed. Boundaries are outlined by dashed lines. Basolateral amygdala (BLA) encompasses LA = lateral amygdala, BA = basal amygdala and BLp = posterior portion of the basolateral amygdala; central amygdala nucleus (CeA) encompasses CeL = lateral portion and CeM = medial portion. Scale bar = 300 µm. (**b**) cFos quantification in the rostral amygdala. Significant effects LA: Kruskal-Wallis one-way ANOVA on ranks (cFos+): P = < 0.001; pairwise Dunn’s test: WT HC vs. WT Ext: P = < 0.05, 2CKO HC vs. 2CKO Ext: P = < 0.01. Significant effects BA: Kruskal-Wallis one-way ANOVA on ranks (cFos+): P = < 0.001; pairwise Dunn’s test: 2CKO HC vs. 2CKO Ext: P = < 0.01. (**c**) cFos quantification in the medial BLA. Significant effects LA: Kruskal-Wallis one-way ANOVA on ranks (cFos+): P = < 0.001; pairwise Dunn’s test: WT HC vs. WT Ext: P = < 0.01, 2CKO HC vs. 2CKO Ext: P = < 0.05. Significant effects BA: Kruskal-Wallis one-way ANOVA on ranks (cFos+): P = < 0.001; pairwise Dunn’s test: WT HC vs. WT Ext: P = < 0.01, 2CKO HC vs. 2CKO Ext: P = < 0.05. (**d**) cFos quantification in the medial CeA. Significant effects CeM: Kruskal-Wallis one-way ANOVA on ranks (cFos+): P = < 0.001; pairwise Dunn’s test: WT HC vs. WT Ext: P = < 0.01, 2CKO HC vs. 2CKO Ext: P = < 0.05. (**e**) cFos quantification in the caudal amygdala. Significant effects LA: Kruskal-Wallis one-way ANOVA on ranks (cFos+): P = < 0.001; pairwise Dunn’s test: 2CKO HC vs. 2CKO Ext: P = < 0.05. Significant effects BA: Kruskal-Wallis one-way ANOVA on ranks (cFos+): P = < 0.001; pairwise Dunn’s test: WT HC vs. WT Ext: P = < 0.01, 2CKO HC vs. 2CKO Ext: P = < 0.05. Significant effects BLp: Kruskal-Wallis one-way ANOVA on ranks (cFos+): P = < 0.001; pairwise Dunn’s test: WT HC vs. WT Ext: P = < 0.05. HC: WT mice (n = 8), 2CKO mice (n = 7); Ext: WT mice (n = 7), 2CKO mice (n = 7). Data are shown as means ± SEM. *P < 0.05, **P < 0.01, ***P < 0.001.

**Supplementary Figure 2.**
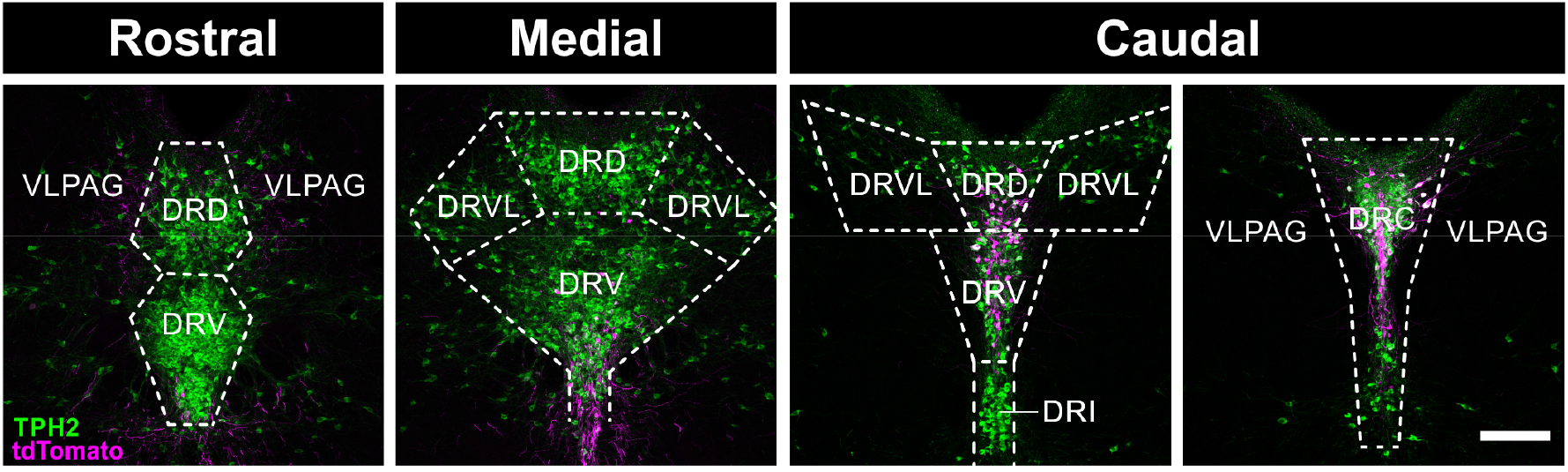
tdTomato expression throughout the DRN. Representative immuno-stained DRN sections of an ePet1-Cre mouse indicate selective tdTomato (magenta) expression in TPH2+ 5-HT cells (green) in the caudal level of the DRN. The pattern of TPH2+ 5-HT cells (green) was used to define the respective level. Boundaries are outlined by dashed lines. DRD = dorsal raphe nucleus, dorsal part; DRV = dorsal raphe nucleus, ventral part; DRI = dorsal raphe nucleus, interfascicular part; DRC = dorsal raphe nucleus, caudal part; DRVL = dorsal raphe nucleus, ventrolateral part; VLPAG = ventrolateral periaqueductal gray. Scale bar = 200 µm.

**Supplementary Figure 3.**
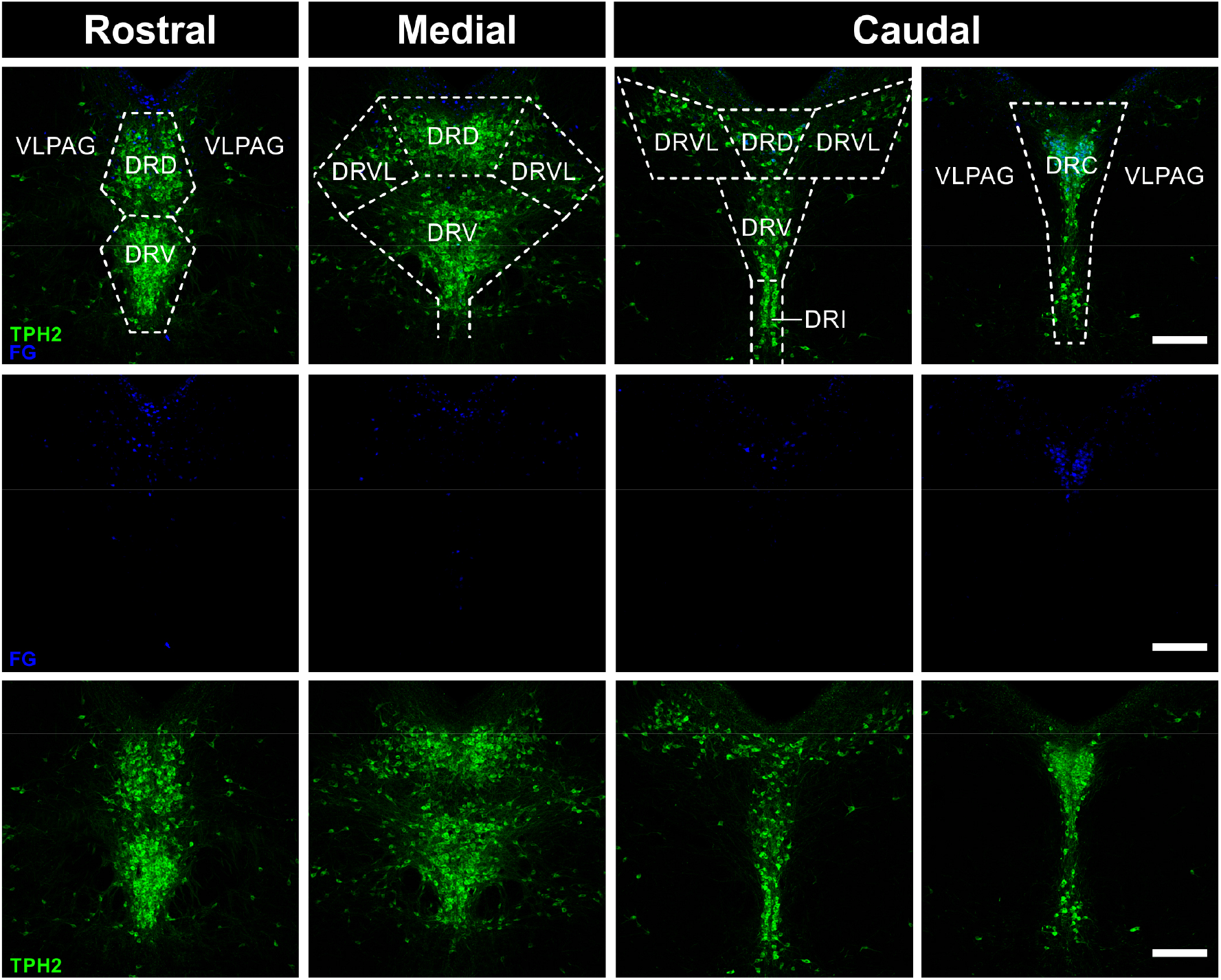
Distribution of retrogradely labeled FG cells in the DRN. Representative immuno-stained DRN sections of a WT mouse injected with 1 % FG into the BNSTad. TPH2+ 5-HT cells (green) in the caudal DRC subregion are densely labeled with FG (blue). Other DRN levels analyzed contain only a few FG labeled TPH2+ 5-HT cells. The pattern of TPH2+ 5-HT cells (green) was used to define the respective level. Boundaries are outlined by dashed lines. DRD = dorsal raphe nucleus, dorsal part; DRV = dorsal raphe nucleus, ventral part; DRI = dorsal raphe nucleus, interfascicular part; DRC = dorsal raphe nucleus, caudal part; DRVL = dorsal raphe nucleus, ventrolateral part; VLPAG = ventrolateral periaqueductal gray. Scale bars = 200 µm.

**Supplementary Figure 4.**
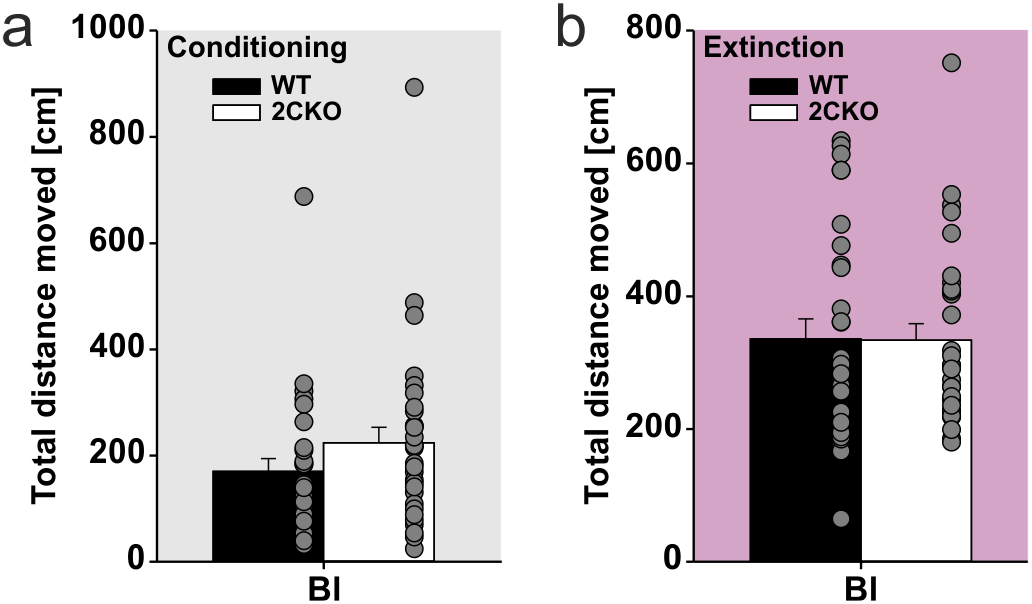
Locomotor activity is not altered in 2CKO mice in the fear conditioning and extinction paradigm. (**a**) Total distance moved during the baseline (Bl) period of the conditioning session (day 2) was similar in both genotypes. (**b**) Total distance moved during the baseline (Bl) period of the extinction session (day 3) was similar in both genotypes. WT mice (n = 29), 2CKO mice (n = 30). Data are shown as means ± SEM.

